# Mother to child blocking of hepatitis B virus and post vaccination serological test in Qinghai province

**DOI:** 10.1101/502658

**Authors:** Ma Xiaojun, Ba Wensheng, Hao Zengping, Li Puren, Zhao Jianhai, Aixitao, Zhu Xianglu, Akezhong, Yang Xueqin, Ma Yanmei, Li Zhongjiu, Li Lianwei, Guan Bingju, Li Xianglan, Zhang Shourong, Li Zhen, Zhao Jinhua, Ma Juan, Samuel So, Zhang Jianmin

## Abstract

**Background:** In 2016, Qinghai Province and the Asian Liver Center of Stanford University in the United States cooperated to carry out a one-year project of hepatitis B mother-to-child blockade and post-immunization serological test in our province. Through this study, we evaluated the current situation of hepatitis B maternal and infant blockade and the effect of maternal and infant blockade in multi-ethnic areas of Qinghai Province.

**Methods:** From May 1, 2016 to April 30, 2017, all pregnant women who gave birth in Qinghai Province were screened for HBsAg. For pregnant women who were screened for HBsAg positive, the medical staff had a detailed understanding of the history and treatment of hepatitis B, and provided scientific nutritional support and guidance. Free hepatitis B immunoglobulin (100 units) injections were given to newborns born to HBsAg-positive mothers within 12 hours after birth, and three doses of hepatitis B vaccination were given. All newborns born to HBsAg-positive mothers were followed up from 1 to 3 months after complete immunization of HBIG and HepB. The County (district) CDC was responsible for rapid fingertip blood screening, screening for HBsAg-positive children, collecting 3ml of venous blood for quantitative detection of HBsAg.

**Results:** During the study period, 61381 pregnant women were hospitalized, 6027 (97.79%) pregnant women were screened for HBsAg. 1912 pregnant women were detected positive for HBsAg, with a positive rate of 3.19%. The HepB vaccine rate was 97.11% in live infants within 24 hours. The vaccination rate of hepatitis B vaccine was 94.78% and the injection rate of HBIG was 97.15% in 12-hour live delivery children of HBsAg positive mothers. A total of 864 newborns born to HBsAg positive mothers were followed up. Fast fingertip blood test was performed 1-3 months after HBIG injection and whole course hepatitis B vaccination. The positive rate of HBsAg was 5.21%. 34 positive serum samples were detected by chemiluminescent immunoassay in the laboratory of provincial CDC. The coincidence rate of HBsAg positive was 82.35%, and that of HBV DNA positive was 79.41%. Fifty-seven children with HBV surface antibody negative were detected and vaccinated with HBV vaccine free of charge in the whole course according to the national immunization program.

**Conclusion:** The infection rate of hepatitis B among women of childbearing age in Qinghai province is 3.19%, and the blocking rate of mother and infant is 94.79%. It is still necessary to strengthen the injection of hepatitis B vaccine and HBIG for positive mothers and children. Keyword: Hepatitis B, Mother to child blocking, Evaluation of the effect

**Author Summary:** This study revealed for the first time the infection rate of hepatitis B virus among women of childbearing age of different nationalities in Qinghai-Tibet Plateau through screening of pregnant women in hospital, blocking of mother and infant, and following-up of serum test after immunization. To evaluate the combined blocking effect of hepatitis B vaccine and HBIG by carrying out mother-to-child blocking and follow-up. The results of this study can provide beneficial reference for the development of hepatitis B virus mother-to-child blocking in minority population in Plateau area.

## 1 Introduction

Hepatitis B is caused by hepatitis B virus, and the burden of disease and financial burden caused by hepatitis B is heavy in China. In countries with high prevalence of HBV, most infections occur in infants, and mother-to-child transmission is the main mode of HBV infection in newborns ^[1–2]^. To prevent mother-to-child transmission of HBV, the American Immune Advisory Committee recommended post-exposure prophylaxis for newborns of HBsAg-positive mothers, i.e., hepatitis B vaccines (HepB) and hepatitis B immunoglobulin (HBIG) within 12 hours after birth, and complete the whole course of three doses of HepB immunization ^[3–5]^. According to the Guidelines for the Prevention and Treatment of Chronic Hepatitis B in China, comprehensive and systematic intervention services should be provided for pregnant and lying-in women to prevent mother-to-child transmission of hepatitis B while carrying out routine maternal health care services. For newborns born to mothers with positive HBsAg, free hepatitis B immunoglobulin (100 units) injections are provided within 12 hours after birth, and three times of hepatitis B vaccination are carried out within 12 hours, 1 month and 6 months of birth according to the requirements of the National Immunization Program. This study is the first time to evaluate the effect of maternal and infant hepatitis B blockade in multi-ethnic areas of Qinghai Province, providing a scientific basis for the prevention and control of hepatitis B.

## 2 method

### 2.1 Investigation area

The survey was carried out in the whole province of Qinghai Province, including Xining, Haidong and six autonomous prefectures (Qinghai Province covers an area of 696,600 square kilometers, with a population of 5.8 million in 2015 and 47% of the minority population). Qinghai Province is located in the western part of China and the northeastern part of the Qinghai-Tibet Plateau. Most of the region belongs to the plateau area. There are many ethnic minorities and the economic development lags behind. According to the geographical and demographic characteristics of Qinghai Province, the whole province is divided into three categories: urban, rural and pastoral areas.

### 2.2 Research object

The subjects were all pregnant and lying-in women who were hospitalized and delivered in Qinghai Province from May 1, 2016 to April 30, 2017. All pregnant and lying-in women with positive HBsAg and their newborns were screened out.

### 2.3 Investigation content

For screening HBsAg-positive pregnant and lying-in women, medical staff should have a detailed understanding of their hepatitis history and treatment, and provide scientific nutritional support and guidance. For newborns born to HBsAg-positive mothers, free hepatitis B immunoglobulin (100 units) injection was provided within 12 hours after birth, Three doses of hepatitis B vaccine (10ug recombinant Hansen yeast) were inoculated. The newborns born to HBsAg-positive mothers were followed up from 1 to 3 months after complete immunization of HBsAg and HBV vaccine. The County (district) CDC was responsible for rapid fingertip blood screening (colloidal gold method), screening for HBsAg-positive persons, collecting 3ml of venous blood from children for quantitative detection of HBsAg. Hepatitis B vaccination and immunoglobulin injection in all neonates during the survey period.

### 2.4 Timely inoculation judgement

The standard HepB1 was inoculated in time: (1) HepB1 was inoculated within 24 hours of neonatal birth. HepB1 timely inoculation rate =HepB1 timely inoculation / live births count 100%.

### 2.5 Statistical analysis

Statistical analysis was carried out with EpiData 3.0 software and Statistical Product and Service Solutions (SPSS) 16.0 software.

### 2.6 Ethics

Told pregnant women in detail the purpose of the survey, And sign the informed consent. blood samples test results back to the mother. The investigation plan was approved by the ethics committee of Qinghai Provincial Center for Disease Control and prevention.

## 3 Results

### 3.1 Basic situation of respondents

During the study period, 61381 pregnant and lying-in women and 57557 live births were investigated.

### 3.2 Prenatal screening rate of HBV markers and HBsAg positive rate

Prenatal maternal serum HBsAg, HBV surface antibody (Anti-HBs), HBVe antigen (HBeAg), HBVe antibody (Anti-HBe) and HBV core antibody (Anti-HBc) were routinely detected by ELISA in hospitals. Domestic reagents were mainly used, accounting for more than 90%. Chemiluminescence was the main detection method, accounting for about 80%, the rest is gold standard. The screening rate of HBV markers was 97.79% and the positive rate of HBsAg was 3.19% in 61381 pregnant and lying-in women. The screening rates of urban, rural and pastoral areas were 99.39%, 95.50% and 92.28% (*x*^2^ = 1715.3, *P* < 0.01), and the positive rates of HBsAg were 2.64%, 4.24% and 4.47% (*x*^2^ = 54.23, *P* < 0.01), respectively. (Table 1 and 2).

**Table 1.**
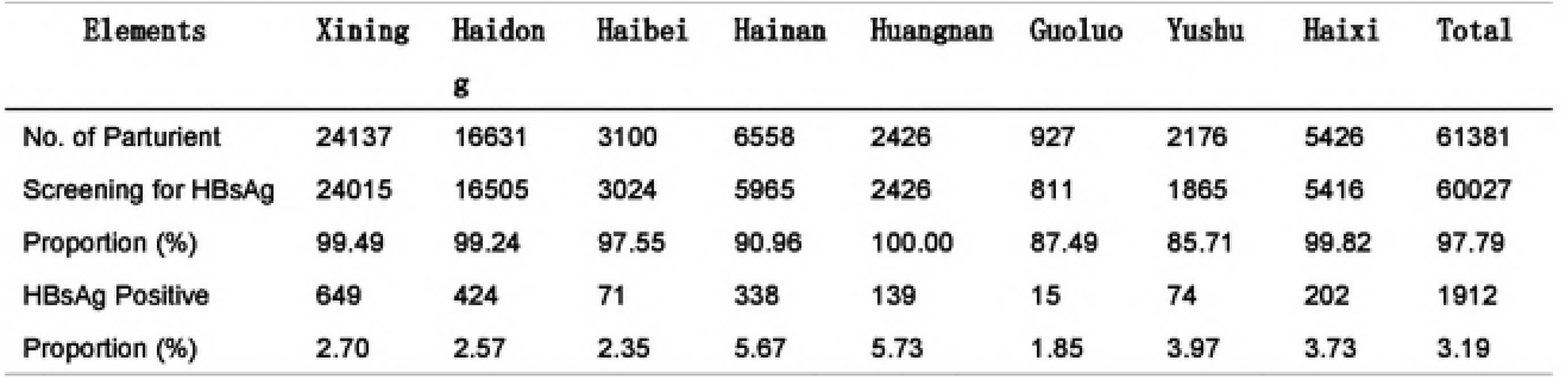
Screening of hepatitis B markers in pregnant and lying in women in Qinghai

**Table 2.**
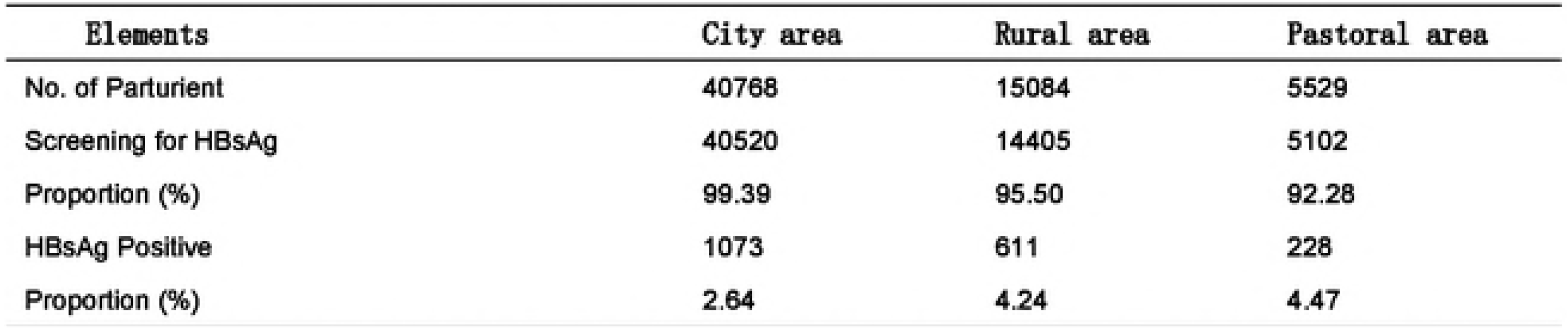
Screening for hepatitis B markers of Parturient in different areas of Qinghai

### 3.3 Neonatal mother to child transmission interruption

#### 3.3.1 Neonatal HepB1 timely vaccination rate

Neonatal HepB1 timely vaccination rate of live children in 24 hours was 97.11%.

#### 3.3.2 The combined inoculation rate of HepB1 and HBIG

There were 1912 positive mothers with hepatitis B surface antigen and 1895 live births. The vaccination rate of hepatitis B vaccine in 12 hours was 94.78%, The rate of HBIG injection in 12 hours was 97.15%, and the rate of combined HBIG and hepatitis B vaccine injection in live-born children of HBsAg-positive mothers was 93.35%.

### 3.4 Maternal and infant blocking effect of newborns born to HBsAg-positive mothers

864 newborns of HBsAg-positive mothers were followed up. HBIG injection and whole course hepatitis B vaccine vaccination were completed. Fast fingertip blood test (colloidal gold method) was performed in 1-3 months. 45 were detected HBsAg (+), and the positive rate was 5.21%. At the provincial level, 34 samples of HBsAg positive infants were received. Thirty-four positive serum samples were detected by chemiluminescent immunoassay in provincial laboratories. 27 of them were HBsAg positive. The coincidence rate of HBsAg positive was 82.35%. 27 cases were positive for HBV DNA, 94.12% of the first dose of hepatitis B vaccine was given in time, 91.18% of the whole course of hepatitis B vaccine and 79.41% of the hepatitis B immunoglobulin(Table 3).

**Table 3.**
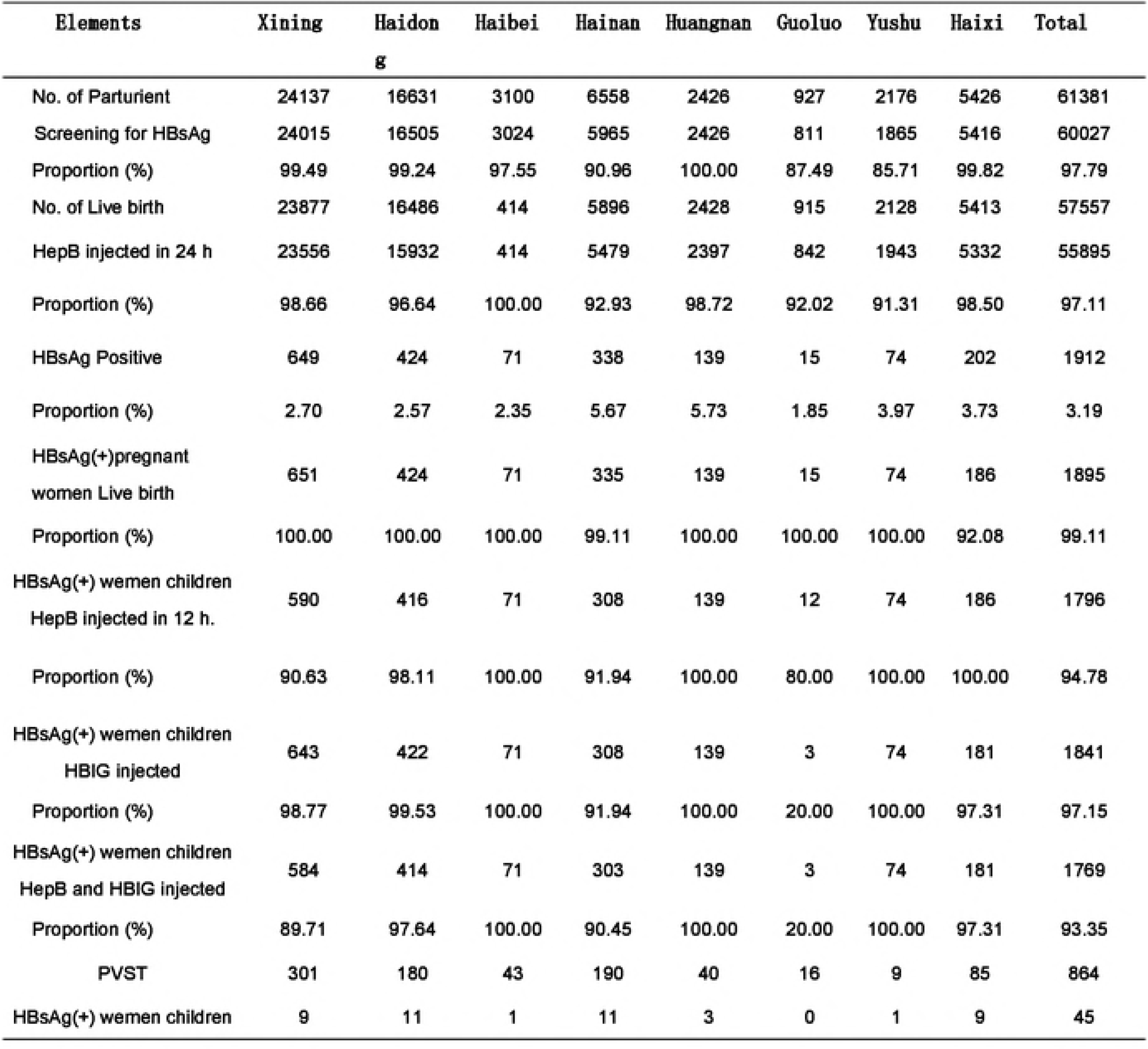
Statistics of hepatitis B markers screening and mother infant blocking work in Qinghai

### 3.5 Post vaccination serological test (PVST) of newborns born to HBsAg-positive mothers

864 newborns born to HBsAg-positive mothers were followed up. Fast fingertip blood tests were performed 1-3 months after HBIG injection and whole course hepatitis B vaccination. 57 children with negative HBV surface antigen and negative HBV surface antibody were detected. According to the national immunization program, they were vaccinated with the whole course of free HBV vaccine again.

## 4 Discussion

Hepatitis B is caused by hepatitis B virus and widely spread worldwide. At present, hepatitis B is still a worldwide public health problem. The burden of disease and economy caused by hepatitis B in China is even heavier. According to the World Health Organization, about 2 billion people worldwide have been infected with HBV, of which 257 million are chronic HBV infections. About 650,000 people die every year from hepatic failure, cirrhosis and primary hepatocellular carcinoma (HCC) caused by HBV infection. HBV infection accounts for 30% and 45% of patients with cirrhosis and HCC worldwide, and 60% and 80% of patients with cirrhosis and HCC in China. In countries with high prevalence of HBV, the positive rate of HBV surface antigen is over 8%. Most infections occur in infants and young children, and mother-to-child transmission is the main mode of HBV infection in newborns ^[1]^. Research data show that, without preventive measures, mothers with HBsAg and HBeAg are both positive, and more than 70%-90% of babies will be infected within one year after birth and become carriers of HBsAg. If mothers with HBsAg positive and HBeAg negative, babies will be infected within one year after birth. 40% were infected. To prevent mother-to-child transmission of HBV, the Advisory Committee on Immunization Implementation of the United States recommended post-exposure prophylaxis for newborns of HBsAg-positive mothers, i.e., hepatitis B vaccines (HepB) and hepatitis B immunoglobulin (HBIG) within 12 hours after birth, and complete the whole course of three doses of HepB immunization. The implementation of maternal and infant blockade can significantly reduce HBV infection, prevent hepatitis B from becoming chronic, and reduce the burden of hepatitis B related diseases. After successful immunization, the immunity against HBV will last at least 20 years ^[3–4]^.

According to the Guidelines for the Prevention and Treatment of Chronic Hepatitis B in China, while carrying out routine prenatal health care services, pregnant and lying-in women should be provided with comprehensive, comprehensive and systematic intervention services to prevent mother-to-child transmission of hepatitis B ^[1]^. For pregnant and lying-in women with positive hepatitis B surface antigen, medical staff should have a detailed understanding of their hepatitis history and treatment, and provide scientific nutritional support and guidance. Free hepatitis B immunoglobulin (100 units) was provided to newborns born to mothers with positive HBsAg. The newborns were injected within 12 hours after birth, and three times of hepatitis B vaccination were carried out within 12 hours, 1 month and 6 months of birth according to the requirements of the national immunization program.

Serological detection of infants born to HBsAg-positive mothers after vaccination refers to the collection of venous blood for detection of HBV serum markers within 12 months after the whole course of immunization with three injections of HBsAg vaccine to determine whether the vaccination has protective effect. Routine post-immunization testing has strong practical significance: first, from the policy level, it can effectively evaluate the mother-infant interruption strategy, timely identify the problems in the implementation of the strategy and adjust them; secondly, from the personal and family level, it can determine whether the vaccination is successful or not, for the vaccination detected by testing. Failure but not infected with HBV susceptible children, early re-vaccination of hepatitis B vaccine, further reduce the risk of HBV infection ^[5–6]^.

WHO Western Pacific region proposed eliminating mother-to-child vertical transmission as the next regional target for hepatitis B prevention and control, considering that in 2017, countries with HBsAg prevalence of children under 5 years of age under 1% could be promoted to carry out post-immunization testing ^[7–9]^. At present, China has not yet requested the comprehensive detection of serum after immunization. Since 2016, all pregnant and lying-in women will be provided with a free examination of hepatitis B surface antigen, and the newborns born to pregnant women with positive hepatitis B surface antigen will be vaccinated with hepatitis B immunoglobulin and hepatitis B vaccine free of charge. The newborns of HBsAg-positive mothers were vaccinated with the first hepatitis B vaccine in time and passive immunization with HBIG. Three doses of hepatitis B vaccine were vaccinated according to the immunization procedure of 0,1 and 6 months. Serum tests were carried out for the infants of HBsAg-positive mothers at the age of 9-12 months. According to the results, parents of children were instructed to carry out hepatitis B vaccine supplementation and reduce the incidence. The risk of HBV infection is ^[10–12]^.

Every year, our province carries out maternal and infant blockade of hepatitis B for newborns, carries out serological tests after immunization for newborns of hepatitis B surface antigen positive mothers and newborns screened for in-patient delivery, and grasps the blocking effect in time. For those who have low or no response to hepatitis B surface antibody after initial immunization, we should take supplementary vaccination measures in time ^[13–15]^. The law was first applied in our province.

The results of the national seroepidemiological survey of hepatitis B in 2006 showed that the HBsAg carrying rate of the population aged 1-59 in China decreased from 9.75% in 1992 to 7.18%. The seroepidemiological survey of hepatitis B among people aged 1-29 in China by CDC in 2014 showed that the detection rates of HBsAg in people aged 1-4, 5-14 and 15-29 were 0.32%, 0.94% and 4.38% respectively. In 2006 and 2014, the detection rate of HBsAg in Qinghai province was lower than the national level ^[16–18]^.

According to the statistics of Qinghai Health and Family Planning Commission, the hospital delivery rates in 2013, 2014 and 2015 were 95.46%, 96.83% and 97.18% respectively, increasing year by year, which provided favorable conditions for screening HBV infection in pregnant women and timely vaccination of HepB1 in newborns. Prenatal screening of HBV infection indicators for pregnant women has important practical significance. The results of screening will help to provide scientific basis for the implementation of mother-to-child transmission interruption and reduce the risk of mother-to-child transmission of hepatitis B ^[19–21]^. In 2011, the former General Office of the Ministry of Health issued the Implementation Plan for the Prevention of Mother-to-Child Transmission of HIV, Syphilis and Hepatitis B. In Qinghai Province, pregnant women were screened for HBV and measures to prevent mother-to-child transmission of hepatitis B were implemented throughout the province. This study shows that the screening rate of HBV markers in pregnant women in Qinghai is 97.79%, which is close to the national average level of ^[22–24]^. Prenatal screening rate of HBV markers can reach 100% in urban areas, but it is still low in rural and pastoral areas. Prenatal screening rate of pregnant women in pastoral areas is 92.28%. Therefore, further awareness needs to be raised in rural and pastoral areas to ensure prenatal screening rate. HepB vaccination in newborns is the most effective method to block mother-to-child transmission of HBV, among which timely inoculation of HepB1 is the most critical ^[25–29]^. This study showed that the timely vaccination rate of HepB1 was 97.11% for newborns delivered in Qinghai Province, 94.78% for newborns born in hospitals by HBsAg-positive mothers, and 93.35% for newborns born in combination with HepB1 and HBIG.

Through the implementation of comprehensive strategy of prevention and treatment of hepatitis B, the awareness of hepatitis B mother-to-child interruption in medical institutions at all levels of the province has been continuously improved. The screening of hepatitis B among pregnant and lying-in women, the vaccination rate of hepatitis B vaccine and the injection rate of HBIG have been continuously improved, which has played an active role in reducing the transmission of hepatitis B. The infection rate of hepatitis B among women of childbearing age in Qinghai Province is 3.19%. For HBsAg positive mother’s children, it is still necessary to strengthen the injection of hepatitis B vaccine and hepatitis B immunoglobulin.

## Acknowledgements

We are grateful to SAMUEL So, Zhang Jianmin. This work benefited from funding support from the Asian Liver Center of Stanford University.

